# Opioids depress breathing through two small brainstem sites

**DOI:** 10.1101/807297

**Authors:** Iris Bachmutsky, Xin Paul Wei, Eszter Kish, Kevin Yackle

**Affiliations:** Department of Physiology, University of California-San Francisco, San Francisco, CA 94143, USA; Neuroscience Graduate Program, University of California-San Francisco, San Francisco, CA 94143, USA; Biomedical Sciences Graduate Program, University of California-San Francisco, San Francisco, CA 94143, USA

## Abstract

The rates of opioid overdose in the United States quadrupled between 1999 and 2017, reaching a staggering 130 deaths per day. This health epidemic demands innovative solutions that require uncovering the key brain areas and cell types mediating the cause of overdose—opioid respiratory depression. Here, we identify two primary changes to breathing after administering opioids. These changes implicate the brainstem’s breathing circuitry which we confirm by locally eliminating the μ-Opiate receptor. We find the critical brain site is the origin of the breathing rhythm, the preBötzinger Complex, and use genetic tools to reveal that just 70-140 neurons in this region are responsible for its sensitivity to opioids. Future characterization of these neurons may lead to novel therapies that prevent respiratory depression while sparing analgesia.

## Introduction

Nearly 400,000 people in the United States died from a drug overdose involving a prescription or illicit opioid between 1999 and 2017 (*1*). This epidemic is not unique to the United States and with the increasing distribution of highly potent synthetic opioids like fentanyl, it has become a global public health emergency (*2*). Death from opioid overdose results from slow and shallow breathing, also known as opioid induced respiratory depression (OIRD, *3*). Like humans, breathing in mice is severely depressed by opioids and this response is eliminated when the μ-Opioid receptor (*Oprm1*) is globally deleted (*4*)*. Oprm1* is broadly expressed, in both the central and peripheral nervous systems, including sites that could modulate breathing such as: the cerebral cortex, brainstem respiratory control centers, primary motor neurons, solitary nucleus, and oxygen sensing afferents (*5,6*). Therefore, either one or multiple sites could be mediating the depressive effects of opioids on breathing.

Indeed, multiple brain regions have been shown to independently slow breathing after local injection of opioid agonists (*6–9*). However, all of these studies fail to definitively demonstrate which of these sites are necessary and sufficient to induce OIRD for three reasons. First, injection of opioid agonists or antagonists into candidate areas modulates μ-opiate receptors on the cell body (post-synaptic) as well as receptors on incoming terminals (pre-synaptic). Second, these studies necessitate anesthetized and reduced animal preparations which alters brain activity in many of the candidate *Oprm1* expressing sites. And third, there is not a ‘gold standard’ definition for how breathing changes in OIRD, enabling each study to use independently designed breathing metrics to measure the impact of their candidate brain site.

To address these limitations, we conducted the first detailed quantitative analysis of OIRD and identify two key changes to the breath that drive the depressive effects of opioids. These two metrics thereafter define OIRD in our study and will serve as a rubric for others to use into the future. We then locally eliminate the μ-opioid receptor in awake mice, disambiguating pre and post-synaptic effects, and use these metrics to define two key brain sites that mediate OIRD. The dominant site is driven by just 140 critical neurons and, importantly, these neurons are not required for opioid-induced analgesia, suggesting a neutral target for developing safer opioids or rescue strategies for opioid overdose.

## Results

Up to now, OIRD has generally been described as a slowing and shallowing of breath (*3*). We therefore felt it was important to more precisely, quantitatively describe the changes in breathing in hopes of elucidating potential mechanisms of respiratory depression. We began by asking whether specific parameters of the breath are affected by opioids. We monitored breathing in awake, behaving mice by whole body plethysmography after intraperitoneal injection (IP) of saline for control and then 20 mg/kg morphine at least 24 hours later (Fig. 1A). Compared to saline, breathing after morphine administration (in normoxia) became much slower and inspiratory airflow decreased, each by 60% (Fig. 1B,C). This culminated in ~50% decrease in overall minute ventilation (MV = tidal volume x respiratory rate, Fig. 1C), demonstrating that 20 mg/kg morphine is, indeed, a suitable dose to model OIRD.

**Fig. 1:**
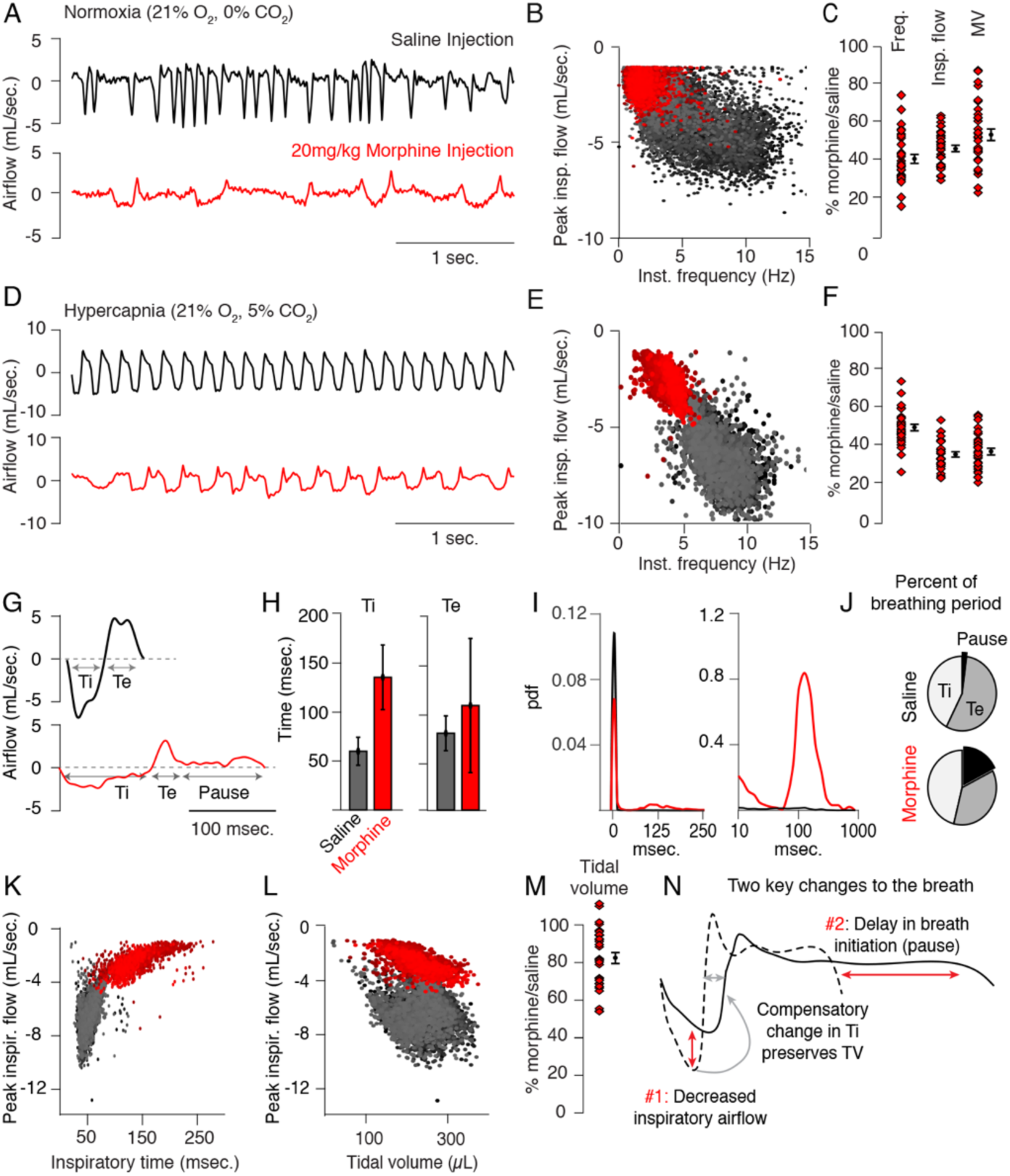
Changes to breathing during opioid-induced respiratory depression. **A,** Representative examples of the breathing airflow (mL/sec.) in normoxia (21% O2) measured by whole body plethysmography. 15 minutes before recordings, animals are intraperitoneally (IP) injected with either saline (black) or 20 mg/kg morphine (red). Morphine recordings are captured 1-7 days after saline. **B**, Scatterplot of instantaneous breathing rate (Hz) versus airflow (mL/sec.) for each breath (dot) taken during the 40-minute recordings. Morphine causes breathing to become slow and less forceful. **C**, Ratio of average breathing parameters after IP injection of morphine-to-saline. Respiratory rate, peak inspiratory airflow, and minute ventilation (MV=tidal volume*rate) for n = 29 animals in normoxia. Red diamond, single animal average. Black diamond, average of all animals. Error bar, standard error of mean (SEM). **D-E**, Representative example of breathing airflow and instantaneous scatter plot (rate vs. airflow) from a 10-minute whole body plethysmography recording of breathing in hypercapnia (21% O2, 5% CO2) to minimize changes in breathing due to differences in behavior after morphine injection. **F**, Ratio of respiratory rate (p-value=1×10^−19^, Cohen’s d=5.96), peak inspiratory airflow (p-value=1×10^−22^, Cohen’s d=6.18), and minute ventilation (p-value=1×10^−20^, Cohen’s d=5.31) after IP injection of morphine-to-saline for n = 29 animals in hypercapnia. **G**, Representative single breath airflow trace for breaths in hypercapnia after saline (black) or morphine (red) IP injection. Hypercapnic saline breaths can be divided into two phases whose durations (msec.) can be measured: inspiration (Ti) and expiration (Te). Hypercapnic morphine breaths have a third phase after expiration where airflow is nearly 0 mL/sec., which we call a pause. **H**, Bar graph of the average length ± standard deviation of Ti and Te for a single representative animal. **I,** Probability density function plot of the pause length in breathing during hypercapnia after saline (black) or morphine (red) IP injection on a numerical (left) and logarithmic scale (right). Note, morphine selectively increases Ti and pause length. **J**, Percent of the average breath period spent in inspiration, expiration, or pause for hypercapnic breaths after saline or morphine injection. **K**, Scatterplot of inspiratory time (msec.) vs. peak inspiratory airflow (mL/sec.) for 10-minutes of hypercapnic breaths after saline (black) or morphine (red) IP injection. As inspiratory time increases, peak inspiratory flow decreases. **L**, Scatterplot of tidal volume (μL.) vs. peak *inspiratory* airflow (mL/sec.) for 10-minutes of hypercapnic breaths after saline (black) or morphine (red) IP injection. Even though peak inspiratory airflow decreases after morphine, tidal volume is preserved due to prolonged Ti. **M**, Ratio of tidal volume after IP injection of morphine-to-saline for n = 29 animals in hypercapnia (p-value=3×10^−6^, Cohen’s d=1.13). **N**, Schematic of the two key morphine induced changes to the breath: decreased inspiratory airflow and pause. Decreased inspiratory airflow prolongs Ti since negative feedback from the lung reflecting breath volume is slower. We interpret the pause as a delay in initiation of the subsequent inspiration.

Breath morphology in normoxia after IP saline versus morphine cannot be directly compared since activity of the mouse is different (exploring vs. sedated, Movie S1-2), which significantly influences the types of breaths taken. This prevented a precise characterization of breath parameters that dictate OIRD. To overcome this, we measured breathing in hypercapnic air (21% O_2_, 5% CO_2_) which normalizes behavior and breathing (Fig. 1D, Movie S3-4). As in normoxia, morphine depressed respiratory rate (by 50%, Fig. 1E,F), peak inspiratory airflow (by 60%, Fig. 1E,F), and minute ventilation (by 60%, Fig. 1F). Hypercapnic breaths after saline exhibited two phases, inspiration and expiration, each lasting about 50 msec. (Fig. 1G,H). After morphine, only the inspiratory phase (measured as inspiratory time, Ti) became substantially longer (Fig. 1G,H). Additionally, hypercapnic breaths showed a new, third phase after expiration that was characterized by little to no airflow (<0.6mL/sec.), and which we therefore define as a pause (Fig. 1G). Such pauses lasted up to several hundred milliseconds (Fig. 1I) and accounted for about one-third of the average breath length (Fig. 1J). Thus, the 50% decrease in respiratory rate after morphine administration is primarily due to prolonging of Ti and pause phases, and the increased prevalence of time spent in pause significantly contributes to the decrease in minute ventilation.

Typically the length of inspiratory time is determined by a stretch-evoked feedback signal from the lung which terminates inspiration (*10*). This reflex is represented by the correlation observed between Ti and peak inspiratory airflow (Fig. 1K). Breaths in morphine still maintain this correlation despite having a longer Ti and decreased inspiratory airflow (Fig. 1K). As a result, morphine breaths have a similar tidal volume (TV) compared to saline control (Fig. 1L,M). In other words, as opioids decrease inspiratory airflow, Ti displays a compensatory increase to preserve TV (Fig. 1N, *11*). In summary, opioids cause only two primary changes to the breath, namely, 1) decreased inspiratory airflow and 2) addition of a pause phase that delays initiation of subsequent breaths (Fig. 1N). These two parameters are controlled by the breathing central pattern generator, the preBötzinger Complex (preBötC), in the brainstem and suggest that this is a key locus affected during OIRD (*12–14*).

Indeed, the preBötC has been proposed to play a key role in OIRD since localized injection of opioids results in respiratory depression and localized naloxone reverses decreased breathing after administration of systemic opioids (*7,15*). However, such experiments fail to distinguish between the action of opioids on presynaptic terminals (*16*) of distant neurons projecting into the preBötC versus direct action on preBötC neurons themselves (Fig. 2A *17,18*). To overcome this, we genetically eliminated the μ-Opiate receptor (*Oprm1*) from preBötC cells exclusively, sparing projecting inputs, by stereotaxic injection of adeno-associated virus constitutively expressing Cre (AAV-Cre) into the preBötC of *Oprm1*(f/f) adult mice (Fig. 2B). To establish a baseline, we first measured breathing after administration of saline and morphine in normoxia and hypercapnia in intact animals, as described above. At least one month after bilateral injection of virus into the preBötC, we then re-analyzed breathing (Fig. 2C). With this protocol, each animal’s unique breathing and OIRD response serves as its own internal control, which is necessary due to the variability in OIRD severity between mice (Fig. 1F). Deletion of *Oprm1* in the preBötC did not affect breathing observed after saline injection (Fig. 2D,E), suggesting that in this context, opiates do not exert an endogenous effect. In contrast, breathing was markedly less depressed by morphine administration (Fig. 2D,E) compared to the intact control state: breaths were twice as fast (3 to 6 Hz, Fig. 2F,H), the peak inspiratory flow was larger (Fig. 2F,I), and pauses were nearly eliminated (Fig. 2G,J). Notably, histological analysis confirmed that AAV-Cre::GFP expression was localized to the preBötC (Fig. S1), and AAV-GFP or tdTomato injected control mice without removal of *Oprm1* showed no change in OIRD compared to the pre-injected control state (Fig. 2H-J), demonstrating that animals do not develop tolerance to opioids within our experimental timeline. Importantly, rescue of OIRD was also specific to breathing since opioids induced analgesia in tail-flick assay after deletion of *Oprm1* in the preBötC (Fig. S2).

**Fig. 2:**
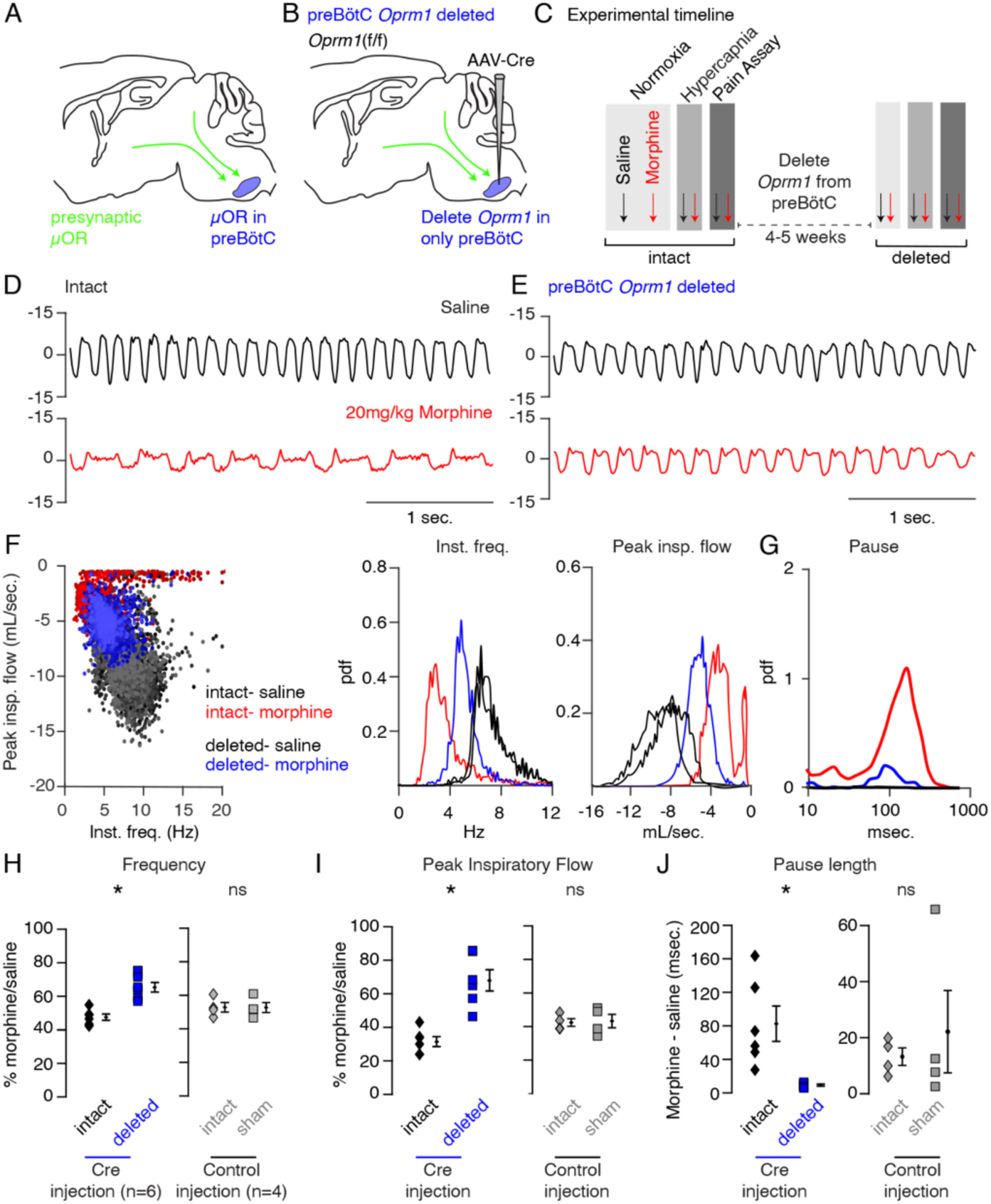
Necessity of preBötC ventrolateral brainstem in opioid-induced respiratory depression. **A,** Schematic of a sagittal section through the adult mouse brain. The μ-Opiate receptor (*Oprm1*) is expressed by a subset of preBötC neurons and is also expressed on presynaptic terminals of some neurons projecting to the preBötC. This confounds the effects observed after localized preBötC injection of opioid or naloxone to investigate its role in OIRD. **B,** To overcome this, we eliminate *Oprm1* only from the preBötC, and not the presynaptic inputs, to define the role of preBötC neurons in OIRD. **C**, Experimental time-course. Breathing is measured 15-minutes after IP injection of saline or 20mg/kg morphine in both normoxia and hypercapnia. Then animals are injected with a constitutive-Cre AAV into the preBötC bilaterally. After several weeks breathing is assayed again as above. In this way, each animal’s breathing before viral injection can serve as its own control breathing. Response to pain is also measured with a tail-flick assay before and after viral injection to ensure that analgesic response is unaffected. **D-E**, Representative examples of the breathing airflow (mL/sec.) in hypercapnia after saline (black) or morphine (red) before (**D**) and after (**E**) Cre-virus injection. **F,** Scatter plot of instantaneous respiratory frequency vs. airflow (ml/sec), as well as probability density function plots of both parameters for a representative animal during hypercapnia after saline (black) or morphine (red, blue) IP injection, before (red) and after (blue) Cre-injection. **G**, Probability density function plot of pause length (msec.) for a representative animal during hypercapnia after IP morphine before (red) and after (blue) Cre-injection, log scale. Prevalence of long duration pauses is greatly reduced. **H-J**, Ratio of average breathing parameters after IP injection of morphine-to-saline. Respiratory rate (**H,** p-value= 0.002, Cohen’s d=3.01), peak inspiratory airflow (**I,** p-value=0.0003, Cohen’s d=3.01), and pause length (**J,** p-value=0.01, Cohen’s d=2.00) for 6 Oprm1(f/f) animals with Cre-virus injected into the preBötC or 4 control animals with reporter-virus injected into the preBötC. For each experiment “intact” values are before viral injection, with “deleted” and “sham” values representing post viral injection conditions in experimental and control animals, respectively. Diamond and square, single animal average. All sham p-values were not significant (>0.2). Black diamond, average of all animals. Error bar, standard error of mean (SEM). * indicates p-value <0.05. ns indicates p-value >0.05.

Although key features of OIRD (inspiratory airflow and pause) were attenuated by preBötC AAV-Cre injection, rescue was incomplete. This could be explained by incomplete *Oprm1* deletion within the preBötC, or participation of another brain site in OIRD. Injection of opioids into the parabrachial (PBN)/Kolliker-Fuse (KF) nucleus can also slow breathing, making it a candidate second site (*8,9*). In fact, the PBN/KF has been proposed to be the key site mediating OIRD (*19,20*). We therefore took a similar approach to test the role of the PBN/KF in OIRD (Fig. 3A). AAV-Cre injection into the PBN/KF (Fig. S3) produced a slight increase in the morphine-evoked respiratory rate (Fig.S4, Fig. 3D,E) and inspiratory airflow (Fig. S4, Fig. 3F,G), but had a more moderate effect than injection into the preBötC.

**Fig. 3:**
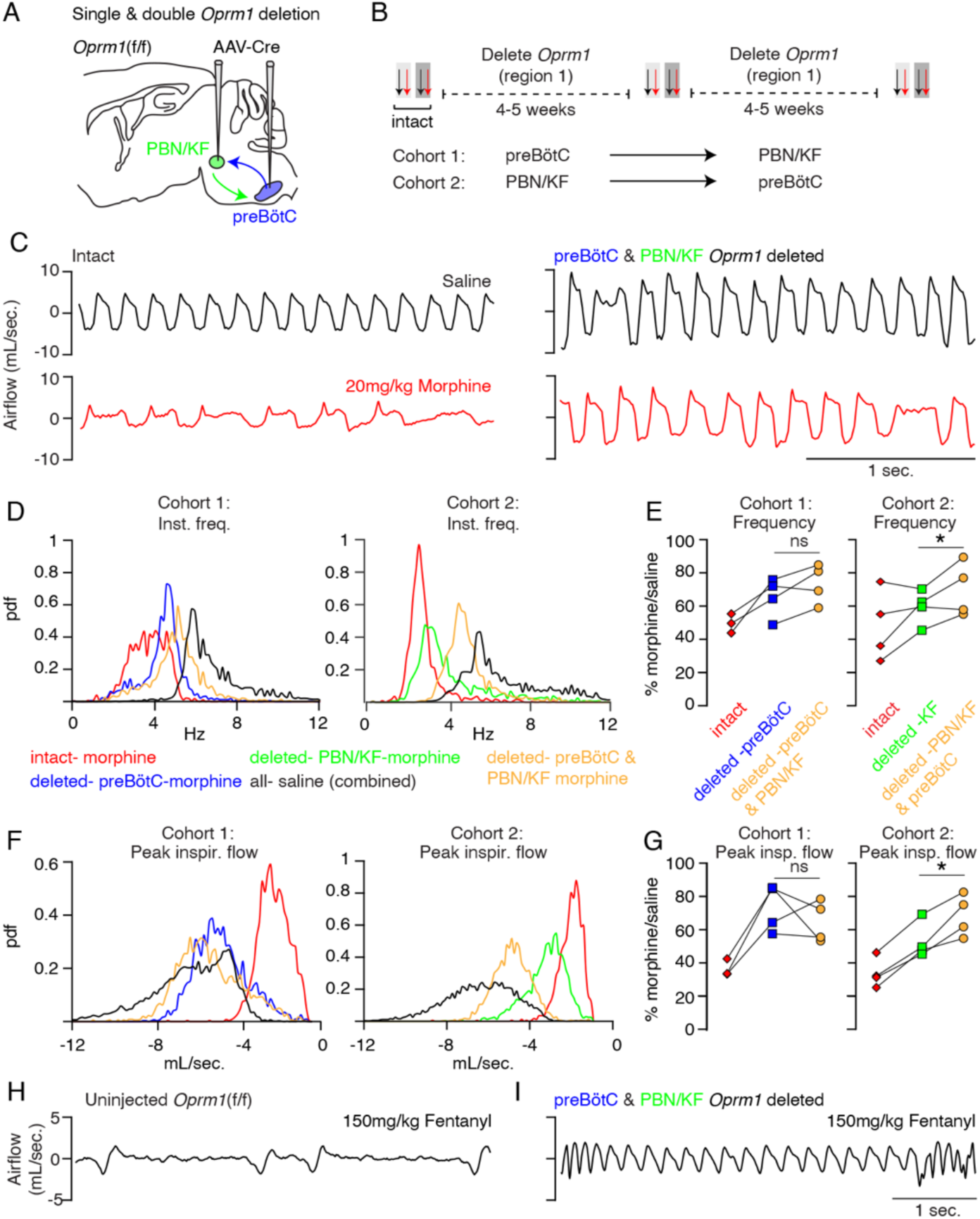
Necessity of Parabrachial/Kolliker Fuse nuclei and preBötC in opioid-induced respiratory depression. **A,** Schematic of a sagittal section through the adult mouse brain showing Cre-viral injection into the preBötC and PBN/KF to measure their individual and combined contribution to OIRD. **B**, Like Fig. 2, breathing was assayed before and after each time Cre-virus was injected. In cohort 1, Cre-virus was first injected into the preBötC and then the PBN/KF and in cohort 2, Cre-virus was injected into the PBN/KF and then preBötC. **C**, Representative examples of the breathing airflow (mL/sec.) in hypercapnia for an animal in cohort 1 after saline (black) or morphine (red) before either viral injection and after both preBötC and PBN/KF Cre-virus injections. **D-E,** Probability density function plot of the instantaneous respiratory frequency for a representative animal from cohorts 1 and 2 (**D**) and ratio of average rate (**E**) after IP injection of morphine-to-saline for 4 animals in cohort 1 and 4 animals in cohort 2 before (pre) and after each Cre-virus injection into the preBötC (blue) or PBN/KF (green). Among the 2 cohorts, n=5 PBN/KF injections were bilateral and n=3 mostly unilateral. Cohort 1: preBötC vs. double p-value=0.07, Cohen’s d=0.69. Cohort 2: PBN/KF vs. double p-value=0.05, Cohen’s d=0.76. **F-G,** Probability density function plot of the peak inspiratory airflow for a representative animal from cohorts 1 and 2 (**F**) and ratio of average peak inspiratory airflow (**G**) after IP injection of morphine-to-saline before (pre) and after each Cre-virus injection into the preBötC (blue) or PBN/KF (green). preBötC Cre-injection has a larger magnitude rescue and after preBötC and PBN/KF injections animals barely have any OIRD phenotype. Cohort 1: preBötC vs. double p-value=0.23, Cohen’s d=0.61. Cohort 2: PBN/KF vs. double p-value=0.01, Cohen’s d=1.32. **H-I,** Representative plethysmography traces in normoxia from a control *Oprm1*(f/f) mouse (N) or a double Cre-injected *Oprm1*(f/f) mouse (preBötC and PBN/KF, O) after IP injection of 150mg/kg fentanyl. * indicates p-value <0.05. ns indicates p-value >0.05.

To determine if the preBötC and PBN/KF can completely account for OIRD (Fig. 3A), we genetically deleted *Oprm1* from the preBötC and then from the PBN/KF (Cohort 1) or vice versa (Cohort 2, Fig. 3B). In either cohort, double deletion breathing after morphine administration looked nearly identical to that of saline control animals (Fig. 3C), with breathing rate and inspiratory airflow depressed by only ~20% compared to saline (Fig. 3D-G). Moreover, changes in breathing after viral injection at the second site appeared additive (Fig. 3D-G) and equivalent to individual preBötC (Fig. 2) or PBN/KF (Fig. S4, Fig. 3D-G) effects for each cognate cohort. To our surprise, rescues also occurred in animals with mostly unilateral PBN/KF AAV-cre injection (Fig. S5), therefore these animals were still included in our double deletion analysis (Fig. 3E,G). Breathing in double-deleted animals was even resilient to super-saturating doses of opioid (150mg/kg fentanyl) that severely slow breathing in control animals (Fig. 3H,I). Taken together, our data are consistent with a model in which both the preBötC and PBN/KF contribute to opioid respiratory depression, with the former being predominant, and together account for OIRD.

Given the relative importance of the preBötC to OIRD, we sought to identify which *Oprm1* expressing cells within this region depress breathing. Single cell transcriptome profiling of the ventral lateral brainstem of P0 mice (Fig. 4A) showed that *Oprm1* (mRNA) is expressed almost exclusively by neurons (Fig. S6) and is remarkably restricted to just 8% of presumed preBötC neurons (Fig. 4B). This alone is interesting, as it suggests that modulation of only a small subset of neurons with the preBötC is enough to significantly impact its ability to generate a rhythm. We also determined that within the preBötC, *Oprm1* (mRNA) was expressed by glycinergic, gabaergic, and glutamatergic neural types alike (Fig. 4C) and therefore *Oprm1* (mRNA) expression is not exclusive to any known rhythmogenic preBötC subpopulation.

**Fig. 4:**
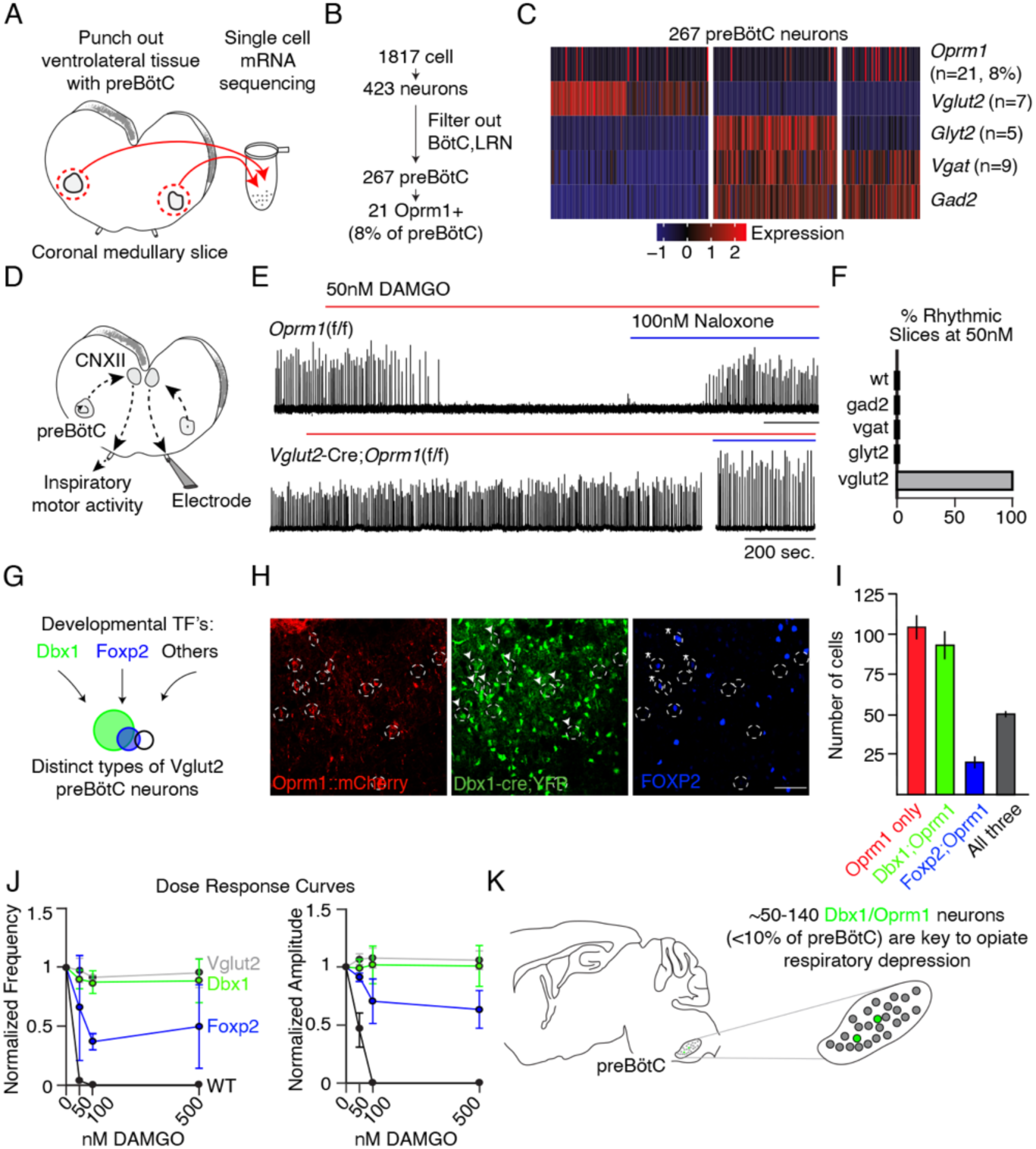
Deletion of μ-Opiate receptor from neural subtypes to define their contribution to opioid depression of preBötC burst rhythm and amplitude. **A,** Schematic of single-cell mRNA sequencing paradigm. Postnatal day 0 (P0) medullary brainstem slices containing the preBötC bilaterally (circled in red) were dissected and isolated for sequencing. **B**, Single cell transcriptome profiling of cells isolated from P0 preBötC. Of 1817 cells isolated, only 267 were presumed preBötC neurons of which only 21 expressed *Oprm1* mRNA. **C**, Heatmap of scaled transcript abundance for *Oprm1* and markers of glutamatergic, gabaergic, and glycinergic preBötC neurons. *Oprm1* expressing cells are both excitatory and inhibitory. **D**, Schematic of extracellular recordings of the preBötC rhythm in P0-4 medullary brainstem slices. The preBötC has input to the hypoglossal motor neurons which form the CN12 rootlet, relaying an inspiratory motor command to the tongue in intact animals. Due to its input from the preBötC, extracellular recording from this rootlet display autonomous rhythmic activity corresponding to *in vitro* respiration (*5*). **E**, Representative recording of bursting activity after application of 50nM DAMGO and 100nM Naloxone. Top: in control (*Oprm1*(f/f)) slices bath application of 50nM DAMGO quickly slowed and decreased the amplitude from baseline bursting. After rhythm cessation, bath application of Naloxone restored the rhythm. Bottom: in *Vglut2*-cre;*Oprm1*(f/f) mice 50nM DAMGO did not stop, or even slow rhythmic activity. **F**, Percent of slices from each genotype with rhythm cessation after 50nM DAMGO application. Control (n=11), *Gad2*-Cre;*Oprm1*(f/f) (n=6), *Vgat*-Cre;*Oprm1*(f/f) (n=2), *Glty2*-Cre;*Oprm1*(f/f) (n=4), and *Vglut2*-Cre;*Oprm1*(f/f) (n=3). **G**, Schematic showing three subpopulations of glutamatergic lineages delineated by transcription factors *Dbx1* and *Foxp2*. *Foxp2* neurons represents a smaller, and overlapping, population of *Dbx1* neurons. **H,** Identification of molecular subtypes of *Oprm1* preBötC excitatory neurons. Sagittal section of the preBötC from a P0 OPRM1::mCherry;*Dbx1*-Cre;Rosa-LSL-YFP mouse immunostained for mCherry (red), YFP (Green) and FOXP2 (Blue). About ~50% of Oprm1 preBötC neurons are glutamatergic/*Dbx1* derived (arrowhead) and of those, ~35% express FOXP2(asterisk). Scale bar, 50μM. **I,** Quantification of the number of preBötC for each molecular subtype identified in **H**. **J**, Dose response curve for the bursting rate and amplitude after bath application of 0, 50, 100, and 500nM DAMGO applied to *Vglut2*-Cre;*Oprm1*(f/f) (gray, n=3), *Dbx1*-Cre;*Oprm1*(f/f) (green, n=3), *Foxp2*-Cre;*Oprm1*(f/f) (blue, n=4) and control (black, n=11) P0-4 preBötC slices. Rate and amplitude for each slice are normalized to baseline. **K**, Schematic summary showing that the key node for opioids to suppress breathing is the preBötC and within this site, elimination of *Oprm1* from just a small subset of those neurons, ~70-140 excitatory neurons, prevents opioid respiratory suppression.

Slices containing the preBötC autonomously generate respiratory-like rhythmic activity *in vitro* which is depressed in both rate and amplitude by bath administration of opioid agonists (*18,21*), similar to opioid effects we observed on breathing *in vivo*. To determine which neural class mediates the depression of preBötC activity, we measured rhythmic bursting activity *in vitro* (Fig. 4D) after selectively genetically deleting *Oprm1* from each neural class. We achieved this deletion by crossing *Oprm1*(f/f) mice with each of the following: *Vglut2*-Cre, *Vgat*-Cre, *Gad2*-Cre, or *Glyt2*-Cre transgenic animals. preBötC slices from control mice (*Oprm1*f/f, f/+, or +/+) burst every 5-10 seconds and this activity was eliminated in 100% of slices by bath application of the selective μ-Opiate receptor agonist [D-Ala^2^, NMe-Phe^4^, Gly-ol^5^]-enkephalin (DAMGO, 50nM), and subsequently rescued by opioid antagonist naloxone (Fig. 4E,F). Strikingly, the bursting rhythm of *Vglut2*-Cre;*Oprm1*(f/f) slices was not slowed by DAMGO, whereas the rhythm in *Gad2*-, *Vgat*-, and *Glyt2*-Cre;*Oprm1*(f/f) slices was entirely eliminated, akin to wild type controls (Fig. 4E,F, Fig. S7). This demonstrates that glutamatergic excitatory neurons, representing ~50% of all preBötC *Oprm1*-expressing neurons and therefore 4% of preBötC neurons, mediate OIRD.

Next, we dissected the glutamatergic *Oprm1* preBötC neurons by two developmental transcription factors, *Dbx1* or *Foxp2* (*22–24*), to determine if a subset can rescue rhythm depression (Fig. 4G). Triple-labeling of *Dbx1*-YFP, OPRM1::mCherry, and FOXP2 protein quantified within a single preBötC revealed three molecular subtypes of P0 *Oprm1* glutamatergic neurons: 92 ± 9 Dbx1, 50 ± 2 Dbx1/FOXP2, and 20 ± 4 FOXP2 (Fig. 4H,I). We selectively eliminated the μ-Opiate receptor in these two lineages (*Dbx1*-Cre;*Oprm1*(f/f) or *Foxp2*-Cre;*Oprm1*(f/f)) and measured preBötC slice activity at increasing concentrations of DAMGO, exceeding the dose necessary to silence the control rhythm (500nM vs. 50nM). Elimination of *Oprm1* from both genotypes was sufficient to rescue the frequency and amplitude of preBötC bursting in DAMGO, and the Dbx1 rescue was comparable to elimination of *Oprm1* from all glutamatergic neurons, while the Foxp2 rescue was substantial, but partial (~50-60%, Fig. 4J). This shows that opioids silence a small cohort (~140) of glutamatergic neurons to depress preBötC activity, and that a molecularly defined subpopulation, about half, can be targeted to rescue these effects.

## Discussion

Here we show that two small brainstem sites are sufficient to rescue opioid induced respiratory depression. Between them, the preBötC is the critical site and we molecularly define ~140 *Oprm1* glutamatergic neurons within that are responsible for this effect. Future study of these neurons will provide the first example of endogenous opiate modulation of breathing. Furthermore, characterization of these neurons and their molecular response to opioids may reveal a strategy for separating respiratory depression from analgesia and therefore enable the development of novel opioids or related compounds that relieve pain without risk of overdose.

### A rubric for studying opioid and other respiratory depressants

We find that although multiple breathing parameters are impacted by opioids, decreased inspiratory airflow and delayed breath initiation, which we term pause, represent the primary changes that result in OIRD. The force and timing of inspiration are ultimately determined by the inspiratory rhythm generator, the preBötC, and focused our initial studies to this site. From here on, these two changes in the breath can act as a ‘gold standard’ for OIRD and should guide future studies charactering and testing novel opioid drugs. Additionally, this workflow can be applied to the analysis of other respiratory depressants.

### Just two small brainstem sites mediate OIRD

Our experimental design allowed us to determine that both the preBötC and KF/PBN have independent and additive rescue of OIRD. Of these two, the preBötC has the larger magnitude rescue of our two core breathing parameters. The combined deletion of μ-Opiate receptor from both sites essentially eliminates OIRD, even to extremely high doses of the potent opioid fentanyl. This suggests that targeting just these two sites is sufficient to rescue opioid respiratory depression. We interpret the small remaining effect of opioids to be due to incomplete transduction of these brain areas but cannot rule out one or more other minor contributing sites.

### Depression of preBötC rhythm by silencing a small glutamatergic subpopulation

The two hallmark changes during OIRD, decreased inspiratory airflow and delayed initiation, perfectly match the opioid induced depression of amplitude and frequency in the preBötC slice. We show that ~140 *Oprm1* glutamatergic preBötC neurons mediate this effect. And surprisingly, half this number, just ~70 glutamatergic neurons are sufficient to rescue opioid depression of the preBötC rhythm (50 Dbx1/FOXP2 and 20 FOXP2 in *Foxp2-*Cre;*Oprm1*(f/f), Fig. S8). Further, given the importance of Dbx1 neurons in respiratory rhythm generation (*22,23*), rescuing *Oprm1* in just ~50/140 Dbx1 neurons (the Foxp2+ subset) may be sufficient to prevent preBötC depression, the smallest number of neurons we propose. This small number is remarkably consistent with the number of Dbx1 neurons that must be lesioned to arrest preBötC activity (*25*). Given the similarity of these effects, we hypothesize that opioids are primarily acting by silencing presynaptic release, effectively removing these neurons from the network. It is profound that such a small number can abruptly halt the respiratory rhythm in a network of more than 1000 neurons. This suggests that either these neurons act as a key population for rhythmogenesis, or that recurrent excitatory networks are exquisitely sensitive to the number of participating cells.

While this manuscript was in preparation a paper appeared that presents some findings similar to ours (*26*).

## Materials and Methods

### Animals

*Oprm1*(f/f) (*27*), *Oprm1*::mCherry (*28*), *Vglut2*-Cre (*29*), *Gad2*-Cre (*30*), *Vgat*-Cre (*29*), *Glyt2*-Cre (*31*), *Dbx1*-Cre (*32*), *Foxp2*-Cre (*33*), Rosa-LSL-YFP (*34*) have been described. Littermates of transgene-containing mice were used as wild type controls. C57Bl/6 mice were used for single cell mRNA sequencing. Mice were housed in a 12-hour light/dark cycle with unrestricted food and water. *Oprm1*(f/f) mice were assigned into experimental and control groups at weaning and given anonymized identities for experimentation and data collection. All animal experiments were performed in accordance with national and institutional guidelines with standard precautions to minimize animal stress and the number of animals used in each experiment.

### Recombinant viruses

All viral procedures followed the Biosafety Guidelines approved by the University of California, San Francisco (UCSF) Institutional Animal Care and Use Program (IACUC) and Institutional Biosafety Committee (IBC). The following viruses were used: AAV5-CMV-Cre-GFP (4.7×10^19^ particles/mL, The Vector Core at the University of North Carolina at Chapel Hill), AAV5-CAG-GFP (1.0×10^13^ particles/mL, The Vector Core at the University of North Carolina at Chapel Hill) or AAV5-CAG-tdtomato (4.3×10^12^ particles/mL, The Vector Core at the University of North Carolina at Chapel Hill).

### Immunostaining

Postnatal day 0-4 brains were dissected in cold PBS, and adult brains were perfused with cold PBS and then 4% paraformaldehyde by intracardiac perfusion. The isolated brains from neonates and adults were then fixed in 4% paraformaldehyde overnight at 4°C and dehydrated in 30% sucrose the next 24 hrs at 4°C. Brains were embedded and frozen in OCT once equilibrated in 30% sucrose. Cryosections (18-25 μM) were washed twice for 5 minutes in 0.1% Tween-20 in PBS, once for 10 minutes in 0.3% Triton-X100 in PBS, and then twice for 5 minutes in 0.1% Tween-20 in PBS. Following wash, sections were blocked for 20 minutes with either 10% goat serum in 0.3% Trition-X100 PBS. Sections were then incubated overnight at 4°C in the appropriate block solution containing primary antibody. Primary antibodies used were: rabbit anti-SST (Peninsula T-4103, 1:500), rabbit anti-Foxp2 (Abcam ab16046, 1:500), chicken anti-GFP (Abcam ab13970, 1:500), rat anti-mCherry (Life Tech. M11217, 1:500), rabbit anti-Lmx1b (ProteinTech Group 18278-1-AP, 1:500). After primary incubation, sections were washed three times for 10 minutes in 0.1% Tween-20 in PBS, then incubated for 1 hour at room temperature or overnight at 4°C in block containing secondary antibody. Secondary antibodies were: goat anti-rat 555 (Lifetech A21434, 1:200), goat anti-chicken 488 (Lifetech A11039, 1:200), goat anti-rabbit 633 (Lifetech 35562, 1:200). After secondary incubation, sections were washed in 0.1% Tween-20 in PBS and mounted in Mowiol with DAPI mounting media to prevent photobleaching. In the instance where Foxp2 and Lmx1b primary antibodies were used on the same, each primary antibody was preincubated with cognate secondary antibody for 10 minutes and then with excess antigen to the secondary antibody before staining sections overnight at 4°C (Lifetech Zenon staining kit).

### Plethysmography, respiratory and behavioral analysis

Adult (8-20 weeks) *Oprm1*(f/f) mice were first administered either IP 100-200μL of saline or morphine (20mg/kg) and placed in an isolated recovery cage for 15 minutes to allow full onset of action of the drug. Individual mice were then monitored in a 450 mL whole animal plethysmography chamber at room temperature (22°C) in 21% O_2_ balanced with N_2_ (normoxia) or 21% O_2_, 5% CO_2_ balanced with N_2_ (hypercapnia). For fentanyl (150mg/kg) onset of action was so fast (<10sec) that animals were placed directly in the plethysmography chamber after administration of drug. Each session (combination of drug and oxygen condition) was separated by at least 24 hours to allow full recovery. Breathing was monitored by plethysmography, and other activity in the chamber monitored by video recording, for 40-minute periods in normoxia and 10-minute periods in hypercapnia. In cases where mice were subject to single or double site AAV injection to delete *Oprm1* or sham controls, breathing was recorded first before viral injection and then again after deletion (or sham) more than 4 weeks later. Breathing traces were collected using EMKA iOX2 software and exported to Matlab for analysis. Each breath was automatically segmented based on airflow crossing zero as well as quality control metrics. Respiratory parameters (e.g. peak inspiratory flow, instantaneous frequency, pause length, tidal volume, etc) for each breath, as well as averages across states, were then calculated. Instantaneous frequency was defined as the inverse of breath duration. Pause length was defined as the expiratory period after airflow dropped below 0.6 ml/sec. Other respiratory parameters were defined by when airflow crosses the value of 0, with positive to negative being inspiration onset and negative to positive being expiration onset.

Due to limitations in breeding, a power calculation was not explicitly performed before our studies. Studies were conducted on all mice generated; six cohorts of animals. After respiration was measured, mice were sacrificed and injection sites were validated before inclusion of the data for further statistical analysis. We used either paired Student’s t-test or Wilcox Rank Sum test to evaluate statistical significance in comparisons of average (across breaths) pre- and post-morphine respiratory parameters (e.g., peak inspiratory flow, instantaneous frequency), based on the normality of their distributions as defined by Shapiro-Wilk test. In comparisons of intact vs. *Oprm1*-deleted or intact vs. Sham conditions the same tests as above were used to evaluate statistical significance between normalized (morphine/saline, or morphine-saline) respiratory parameters. Normality in this case was determined by Shapiro-Wilk test on the normalized respiratory parameters. All statistics were performed using the publicly available Excel package: Real Statistics Functions.

### Tail flick assays

Mice were injected with saline (control trials) or 20mg/kg morphine. 15-minutes later mice were put into a restraining wire mesh with the tail exposed. One-third of the tail was dipped into a 48-50°C water bath and time was measured for the tail to flick. Immediately after the flick, the tail was removed from the bath. If the tail did not flick within 10-seconds, then the tail was removed. The procedure was video recorded so time to response could be quantified post-hoc. Each mouse was recorded for two saline and two morphine trials.

### Stereotaxic injection

Stereotaxic injections were performed in mice anaesthetized by isoflurane. Coordinates used for the preBötC were: −6.75 mm posterior, −5.05 mm ventral from surface, ±1.3 mm lateral from bregma. Coordinates used for the PBN/KF were: −5.05 mm posterior, −3.7 ventral from surface, ±1.7 lateral from bregma. Injection sites were confirmed by the restricted expression of Cre-GFP, GFP, or tdTomato in the anatomically defined Parabrachial/Kolliker-Fuse (*35*) and preBötC (*12,13*) areas. In the case of preBötC injections, anatomical location of injection site was also confirmed by localization with Somatostatin antibody staining (*36*). After injection of the virus, mice recovered for at least 3-4 weeks before breathing metrics were recorded again. In a subset of animals, mice were then subject to a second site deletion of the complementary brain area, ie. preBötC and then from the PBN/KF (Cohort 1) or vice versa (Cohort 2). These mice were then allowed to recover for another period of at least 3-4 weeks, after which a third set of breathing metrics were recorded.

### Slice electrophysiology

Rhythmic 550 to 650μm-thick transverse medullary slices which contain the preBötC and cranial nerve XII (XIIn) from neonatal *Oprm*(f/f), *Oprm*(f/f);*Vglut2*-Cre+/−, *Oprm1*(f/f);*Gad2*-Cre+/−, *Oprm1*(f/f);*Glyt2*-Cre+/−, *Oprm1*(f/f;)*Vgat*-Cre+/−, *Oprm1*(f/f);*Dbx1*-Cre+/−, *Oprm1*(f/f);*Foxp2*-Cre+/− (P0-5) were prepared as described (*37*). Briefly, slices were cut in ACSF containing (in mM): 124 NaCl, 3 KCl, 1.5 CaCl_2_, 1 MgSO_4_, 25 NaHCO_3_, 0.5 NaH_2_PO_4_, and 30 D-glucose, equilibrated with 95% O_2_ and 5% CO_2_ (4°C, pH=7.4). The rostral portion of the slice was taken once the compact nucleus ambiguus was visualized. The dorsal side of each slice containing the closing of the 4^th^ ventricle. For recordings, slices were incubated with ACSF from above and the extracellular K^+^ was raised to 9 mM and temperature to 27°C. Slices equilibrated for 20 min before experiments were started. The preBötC neural activity was recorded from either XIIn rootlet or as population activity directly from the XII motor nucleus using suction electrodes. Activity was recorded with a MultiClamp700A or B using pClamp9 at 10000 Hz and low/high pass filtered at 3/400Hz. After equilibration, 20min. of baseline activity was collected and then increasing concentrations of DAMGO (ab120674) were bath applied (20nM, 50nM, 100nM, 500nM). Activity was recorded for 20min. after each DAMGO application. After the rhythm was eliminated or 500nM DAMGO was reached, 100nM Naloxone (Sigma Aldrich N7758) was bath applied to demonstrate slice viability. Rhythmic activity was normalized to the first control recording for dose response curves.

### Single Cell mRNA Sequencing and analysis

650 μm-thick medullary slices containing the preBötC were prepared from 10 P0 mice C57Bl/6 mice as described above. The preBötC and surrounding tissue was punched out of each slice with a P200 pipette tip and incubated in bubbled ACSF containing 1mg/ml pronase for 30 minutes at 37°C with intermittent movement. Digested tissue was centrifuged at 800rpm for 1 minute, and the supernatant was discarded and replaced with 1% FBS in bubbled ACSF. The cell suspension was triturated serially with fire-polished pipettes with ~600μm, ~300μm and ~150μm diameter. The cells were filtered using a 40-μm cell strainer (Falcon 352340). DAPI was added to a final concentration of 1μg/mL. The cell suspension was FACS sorted on a BD FACSAriaII for living (DAPI negative) single cells. The cells were centrifuged at 300g for 5 minutes and resuspended in 30 μL 0.04% BSA in PBS. The library was prepared using the 10x Genomics Chromium™ Single Cell 3′ Library and Gel Bead Kit v2 (1206267) and according to manufacturer’s instructions by the Gladstone genomics core. The final libraries were sequenced on HiSeq 4000.

For analysis, sequencing reads were processed using the 10x Genomics Cell Ranger v.2.01 pipeline. A total of 1860 cells were sequenced. Further analysis was performed using Seurat v2.3. Cells with less than 200 genes were removed from the dataset. Data was LogNormalized and scaled at 1e4. Highly variable genes were identified and used for principal component analysis. 25 principal components were used for unsupervised clustering using the FindCluster function. 12 clusters were identified at a resolution of 1.0, displayed in Fig. S5. FindAllMarkers and violin plots of known cell type markers were used to identify each cluster.

## Acknowledgments

We thank Matthew Collie (currently Harvard University) for his scientific input throughout the project and experimental contributions to the slice electrophysiology. We thank Drs. David Julius and Roger Nicoll (University of California, San Francisco) for input and revision of the manuscript. We thank Dr. Massimo Scanziani (University of California, San Francisco) for mentorship, discussion, and revision of the manuscript. We thank Adelae Durand (Yackle lab, University of California, San Francisco) for data collection and her revision of the manuscript.

## Author Contributions

I.B. and E.K. performed mouse stereotaxic injections. I.B. and E.K. conducted breathing plethysmography experiments and analgesia assays. I.B., P.W., and K.Y. conducted slice electrophysiology experiments. P.W. performed single cell sequencing and analysis. I.B. and K.Y. performed histological analysis. I.B. performed all analysis on breathing plethysmography and slice electrophysiology data. I.B. and K.Y. conceived experiments, interpreted data and wrote the manuscript. All authors edited the manuscript.

## Supplementary Materials

**Fig. S1:**
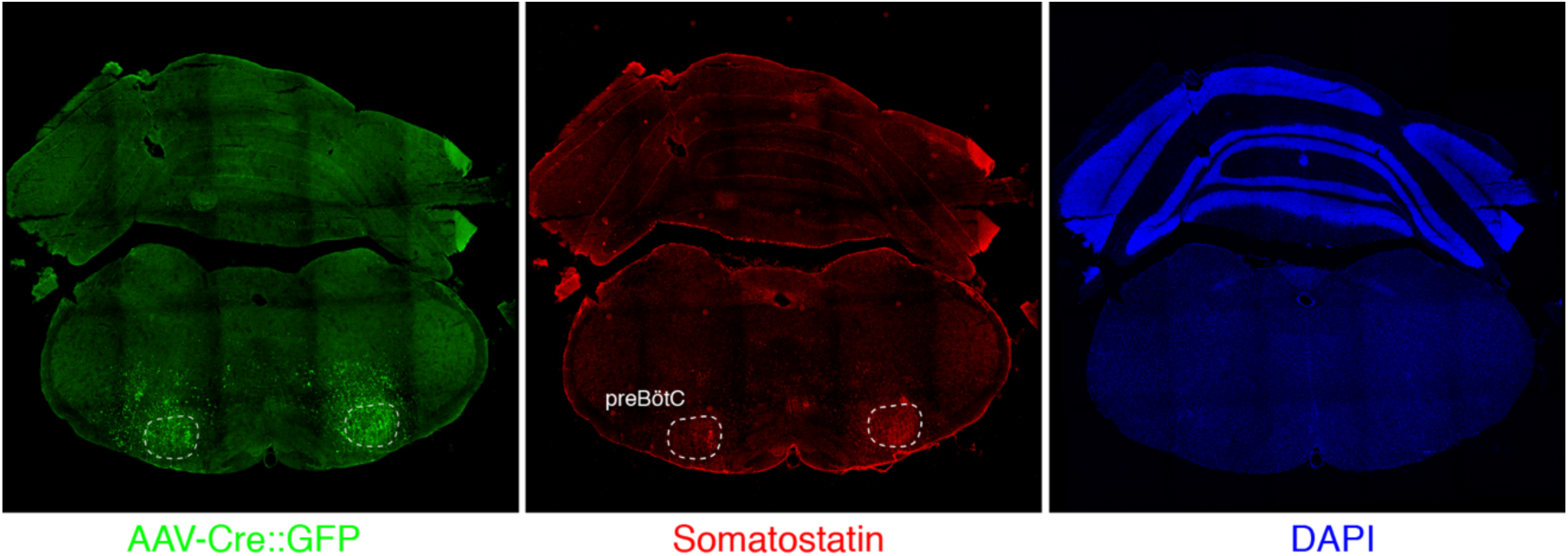
Epression of GFP protein from AAV-Cre::GFP after bilateral preBötC injection. GFP protein (green) and Somatostatin neuropeptide (red) expression in a 25μM coronal section of the brainstem from an adult *Oprm1*(f/f) mouse injected bilaterally with AAV-Cre::GFP into the preBötC. Somatostatin expression demarcates the preBötC (dashed white circles, *34*). This validates the specific targeting of AAV-Cre, and thus *Oprm1* deletion, to the preBötC.

**Fig. S2:**
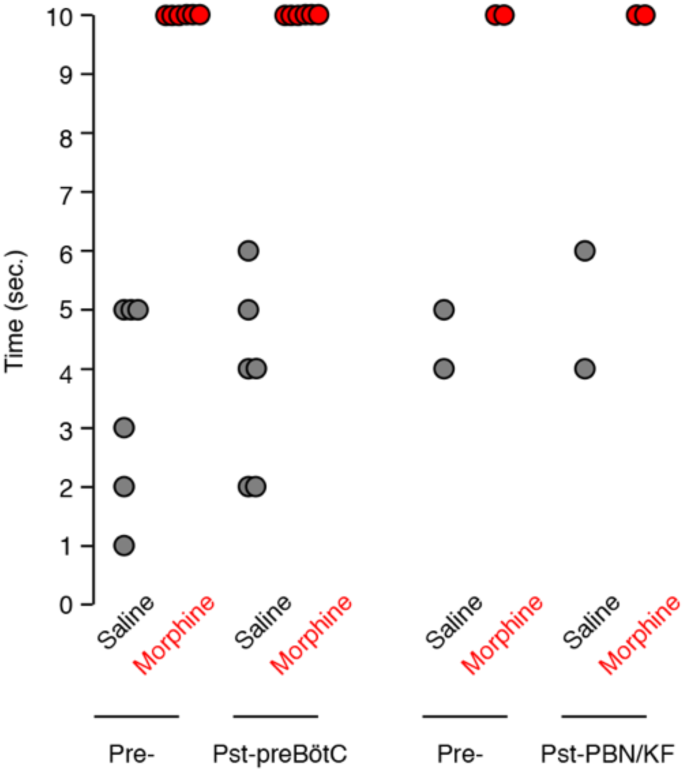
Tail flick response before and after bilateral preBötC or PBN/KF injection. The tail flick response time was measured 15-minutes after IP saline (gray) or 20mg/kg morphine (red) injection. Each animal was assayed twice before (Pre-) and twice after (Pst-) bilateral AAV-Cre::GFP injection into either the preBötC or PBN/KF of adult *Oprm1*(f/f) mice. Dot, time for single assay of one animal. Note, *Oprm1* in the preBötC and PBN/KF is not required for morphine’s ability to blunt the tail flick response.

**Fig. S3:**
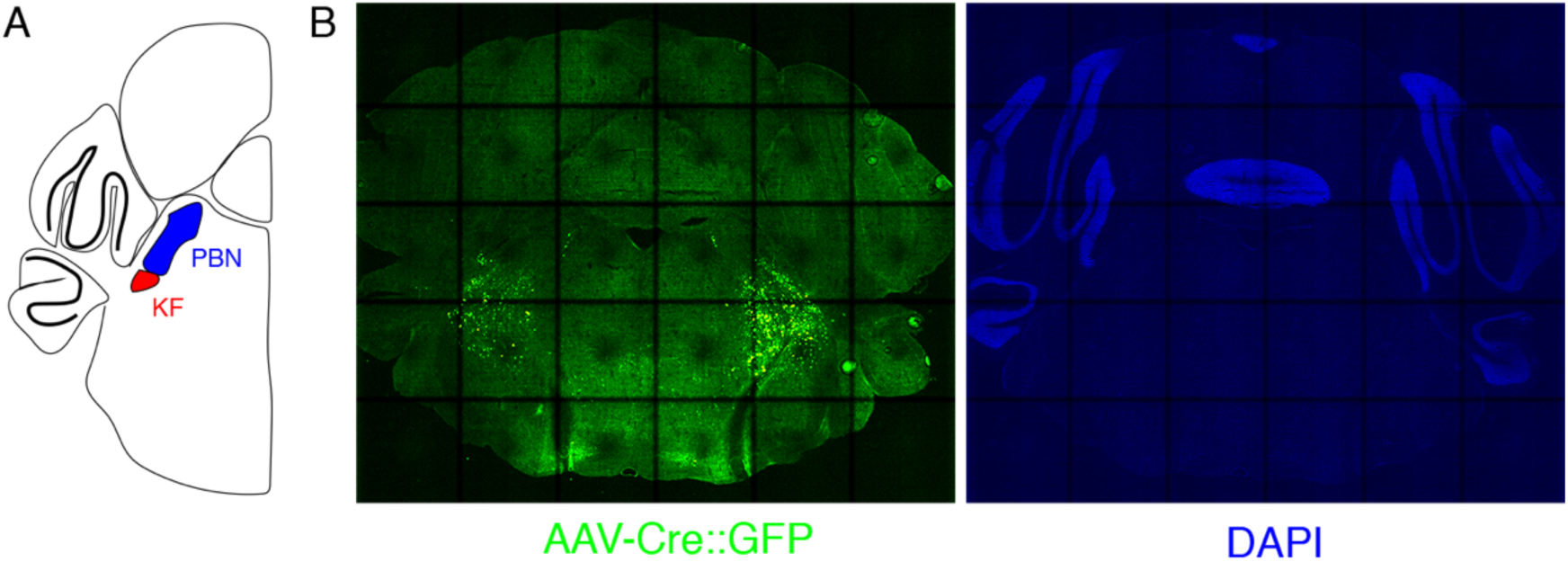
Expression of GFP protein from AAV-Cre::GFP after bilateral PBN/KF injection. **A**, Schematic of a coronal section through the brainstem at the position of the PBN/KF. The PBN/KF is in the dorsal brainstem near the cerebellar peduncle. Half the brain is represented here. **B**, GFP protein (green) expression in a 25μM coronal section of the brainstem from an adult *Oprm1*(f/f) mouse injected bilaterally with AAV-Cre::GFP into the PBN/KF. This anatomical localization of GFP validates the specific targeting of AAV-Cre, and thus *Oprm1* deletion, to the PBN/KF.

**Fig. S4:**
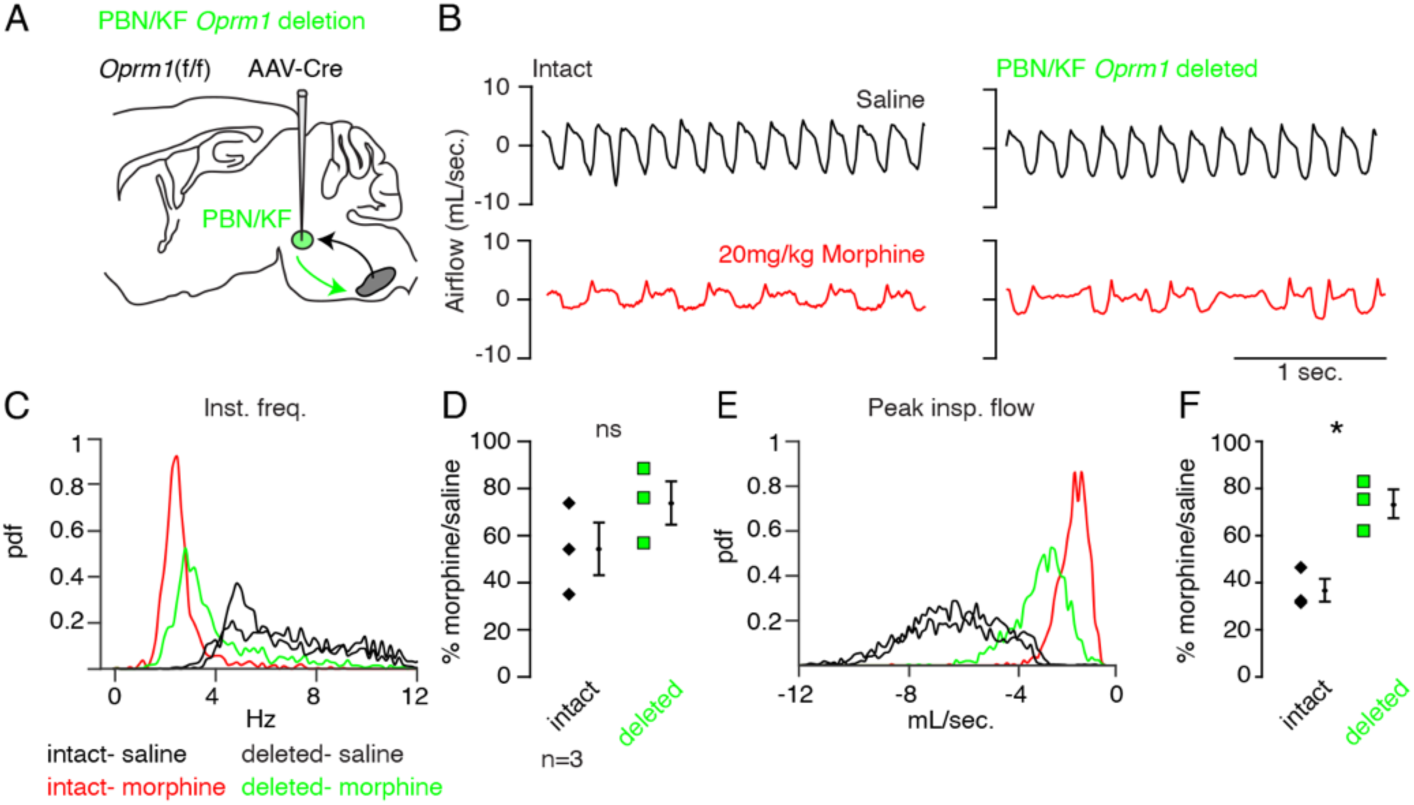
Necessity of Parabrachial/Kolliker Fuse nuclei in opioid-induced respiratory depression. **A,** Schematic of a sagittal section through the adult mouse brain. *Oprm1* is expressed by neurons within the pontine Parabrachial/Kolliker Fuse (PBN/KF) which has modulatory input into the preBötC (*38,39*) and is a proposed region for respiratory depression during OIRD (*14,15*). As in Fig. 2, breathing was assayed before and after Cre-virus injection into the PBN/KF. **B**, Representative examples of the breathing airflow (mL/sec.) in hypercapnia after saline (black) or morphine (red) before and after Cre-virus injection. **C-D,** Probability density function plot of the instantaneous respiratory frequency for a representative animal (**C**) and ratio of average rate (**D,** p-value=0.2, Cohen’s d=0.62) after IP injection of morphine-to-saline for 3 animals before (pre) and after (post) Cre-virus injection into the PBN/KF. **E-F,** Probability density function plot of the peak inspiratory airflow for a representative animal (**E**) and ratio of average peak inspiratory airflow (**F,** p-value=0.01, Cohen’s d=1.68) after IP injection of morphine-to-saline for 3 animals before (pre) and after (post) Cre-virus injection into the PBN/KF. * indicates p-value <0.05. ns indicates p-value >0.05.

**Fig. S5:**
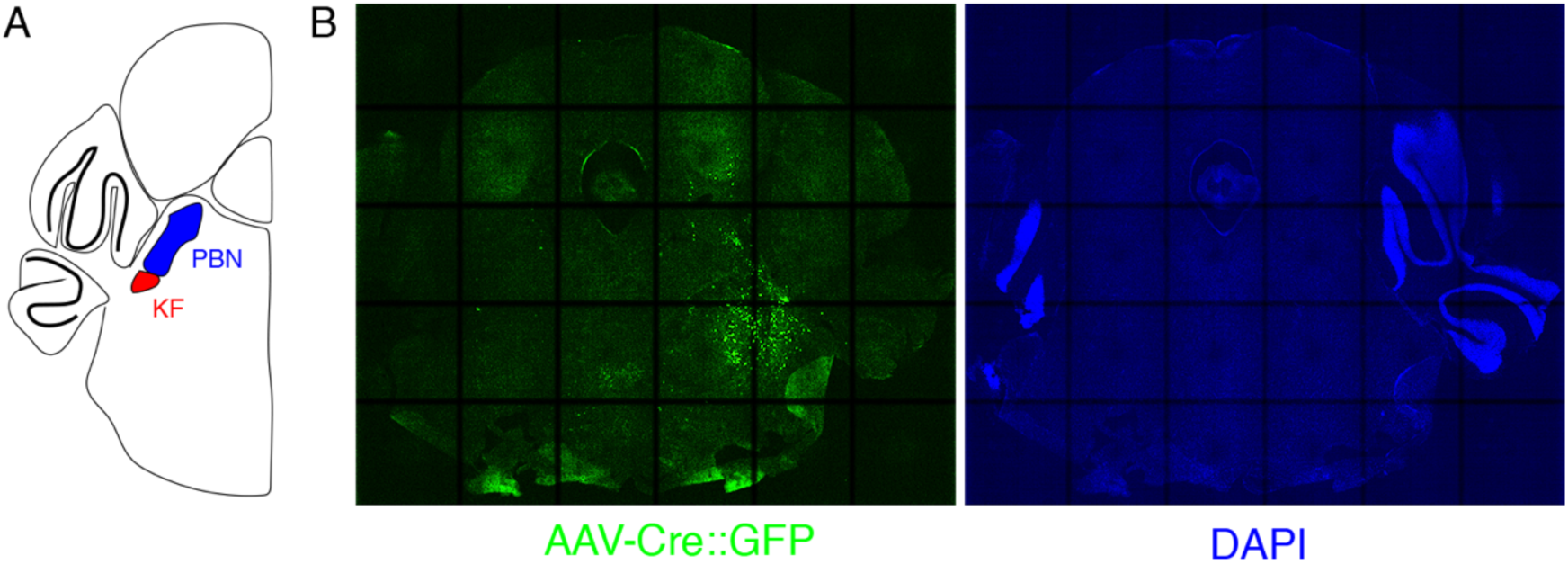
Expression of GFP protein from AAV-Cre::GFP after mostly unilateral PBN/KF injection. **A**, Schematic of a coronal section through the brainstem at the position of the PBN/KF. The PBN/KF is in the dorsal brainstem near the cerebellar peduncle. Half the brain is represented here. **B**, GFP protein (green) expression in a 25μM coronal section of the brainstem from an adult *Oprm1*(f/f) mouse injected bilaterally with AAV-Cre::GFP into the PBN/KF. In this instance, GFP expression is robustly seen on right side and few cells are labeled on the left. This localization of GFP suggests targeting of AAV-Cre, and thus *Oprm1* deletion, is only in to one PBN/KF.

**Fig. S6:**
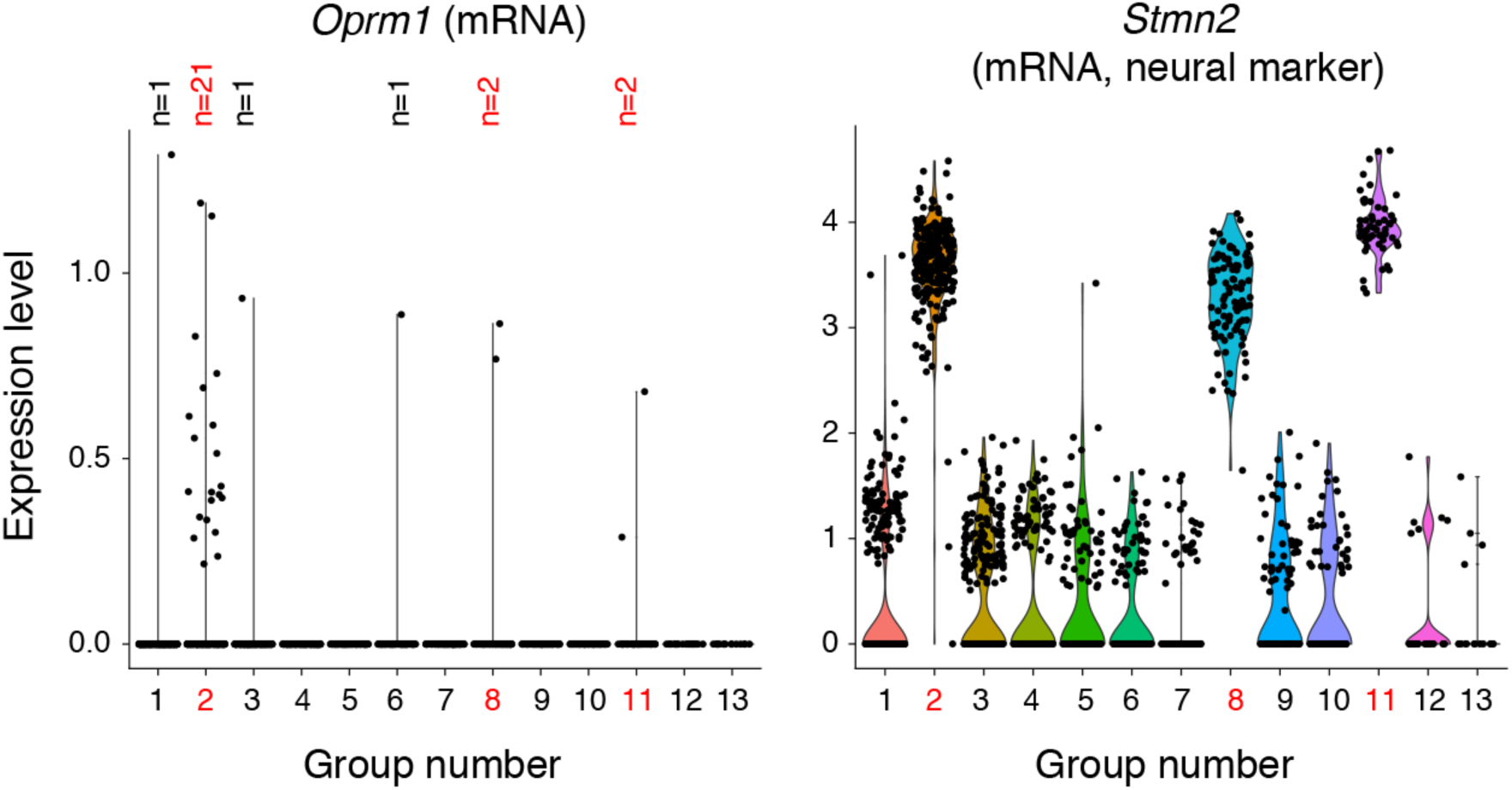
Single cell transcriptome profiling of ventrolateral brainstem neurons. 1817 single cells were isolated from the ventrolateral medulla of the P0 mouse. The transcriptome was sequenced with 10X genomics technology and cells were clustered by principle components analysis into 13 groups. 3 groups (2, 8, and 11, red numbers) expressed neural markers like *Stmn2* (mRNA) shown here. Group 2 was a mixture of excitatory and inhibitory neurons and is the presumed preBötC cluster due to expression of *Sst*, *Tacr1*, and *Cdh9* (mRNA, not shown). Group 8 is a BötC cluster based on expression of *Gad1* and *Neurod6* (mRNA, not shown). Group 11 is a lateral reticular nucleus cluster based on expression of *Zic1* and *Barhl1* (mRNA, not shown). All other groups were oligodendrocytes, oligodendrocyte precursor cells, astrocytes, microglia and endothelial cells. The only group with enriched *Oprm1* (mRNA) expression is the preBötC group 2. Within this group only 21/267 neurons express *Oprm1* (mRNA). The single cell in group 1, 3, 6 (which are non-neural) that express *Oprm1* (mRNA) are presumed contaminants.

**Fig. S7:**
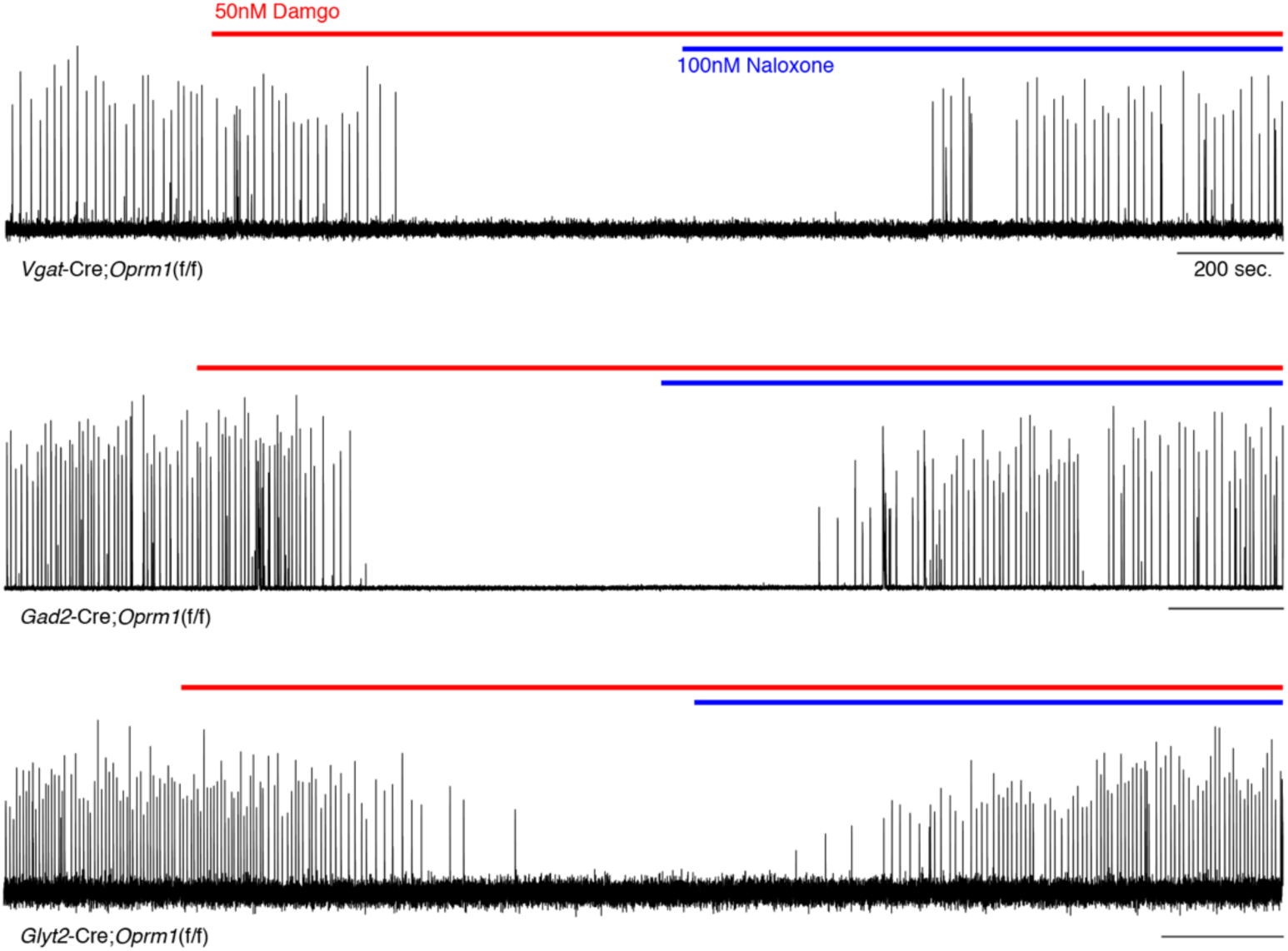
preBötC slice activity in 50nM DAMGO after deletion of *Oprm1* from inhibitory neural types. Representative recording of bursting activity at baseline, after application of 50nM DAMGO (red line), and 100nM Naloxone (blue line). Top: *Vgat*-Cre;*Oprm1*(f/f) targeting GABAergic neurons (n=2). Middle: *Gad2*-Cre;*Oprm1*(f/f) targeting GABAergic neurons (n=6). Bottom: *Glyt2*-Cre;*Oprm1*(f/f) targeting Glycinergic neurons (n=4).

**Fig. S8:**
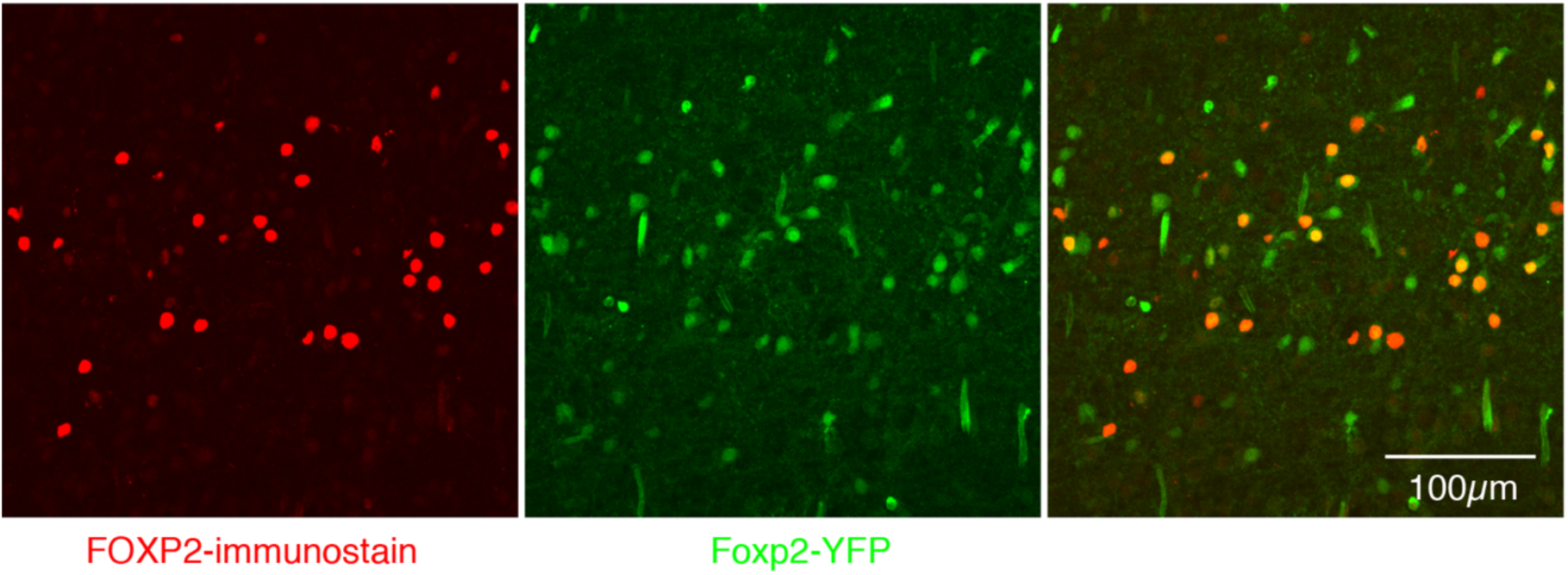
Expression of FOXP2 protein and in Foxp2-derived cells within the preBötC. Nearly perfect colocalization of FOXP2 protein (red) and YFP (green) in a 25μM sagittal section of the preBötC from a P0 *Foxp2*-Cre;Rosa-LSL-YFP mouse. This validates the use of *Foxp2*-Cre to remove *Oprm1* from Foxp2-derived neurons. Scale bar, 100μM.

**Movie S1.**

30 second representative movie during a baseline recording of breathing in normoxia and video documentation of behaviors performed in plethysmography chamber 20 minutes after IP injection of saline. Under this condition mice spend most of their time exploring and sniffing (5-19 seconds in video) or grooming themselves (19-35 seconds).

**Movie S2.**

30 second representative movie during a morphine recording of breathing in normoxia and video documentation of behaviors performed in plethysmography chamber 20 minutes after IP injection of 20mg/kg morphine. Under this condition mice spend most of their time calm (5-24 seconds in video) and sedated (24-35 seconds) or grooming themselves (not shown). This is unlike their behavior after IP saline.

**Movie S3.**

30 second representative movie during a baseline recording of breathing in hypercapnia and video documentation of behaviors performed in plethysmography chamber 20 minutes after IP injection of saline. Under this condition mice are mostly not exploring and instead immobile and breathing heavily.

**Movie S4.**

30 second representative movie during a morphine recording of breathing in hypercapnia and video documentation of behaviors performed in plethysmography chamber 20 minutes after IP injection of 20mg/kg morphine. Under this condition mice behave similar to control saline: mostly immobile and breathing heavily.

